# CCR2 limits inflammatory functions of CD8 TRM cells that impair recognition memory during recovery from WNV encephalitis

**DOI:** 10.1101/2024.09.17.613307

**Authors:** Shenjian Ai, Artem Arutyunov, Joshua Liu, Jeremy D. Hill, Xiaoping Jiang, Robyn S. Klein

## Abstract

Central nervous system (CNS) resident memory CD8 T cells (T_RM_) that express IFN-γ contribute to neurodegenerative processes, including synapse loss, leading to memory impairments. Here, we show that CCR2 signalling in CD8 T_RM_ that persist within the hippocampus after recovery from CNS infection with West Nile virus (WNV) significantly prevents the development of memory impairments. Using CCR2-deficient mice, we determined that CCR2 expression is not essential for CNS T cell recruitment or virologic control during acute WNV infection. However, transcriptomic analyses of forebrain CCR2^+^ versus CCR2^-^ CD8 T_RM_ during WNV recovery reveal that CCR2 signalling significantly regulates hippocampal CD8 T_RM_ phenotype and function via extrinsic and intrinsic effects, decreasing the expression of CD103 and granzyme A and IFN-γ, respectively. Consistent with this, WNV-recovered *Cd8a*^cre^*Ccr2*^fl/fl^ mice exhibit decreased recognition memory. Our findings highlight a neuroprotective role for CCR2 in limiting CD8 T cell-mediated neuroinflammation and cognitive deficits, providing insights into potential therapeutic targets for CNS infections.

## Introduction

T cells, a critical component of the adaptive immune system, play a pivotal role in immune surveillance and defense mechanisms within the central nervous system (CNS) with active roles in CNS homeostasis^1^ and virologic control^2,3^. However, recent studies indicate resident memory CD8 T (T_RM_) cells may also contribute to neurodegenerative processes during recovery from viral infections, including West Nile virus^4^ (WNV) and severe-acute-respiratory-syndrome-related coronavirus (SARS-CoV)-2^5^, and in patients with Alzheimer’s diseases^6,7^. Thus, significant advances in understanding the roles of CD8 T_RM_ cells in the CNS, especially their protective versus detrimental effects in different diseases, require a more comprehensive investigation into their fundamental biology to pave the way for novel therapeutic interventions targeting T cell-mediated processes in various CNS diseases^8^.

WNV is a mosquito-borne arbovirus from the Flaviviridae family, which also includes Zika (ZIKV) and Japanese encephalitis (JEV) viruses. In adults, WNV targets fully differentiated neurons in the CNS^9^ and causes severe long-term cognitive and memory impairment in survivors of WNV fever or neuroinvasive disease (WNND)^10^. During acute WNV infection, the blood-brain barrier is disrupted, accompanied by leukocyte infiltration into the CNS that may permanently alter its cellular milieu^11^. In prior studies, we established a murine model to study the underlying mechanisms of neurological complications during flavivirus encephalitis recovery using the mutant strain, WNV-NS5-E218A, which contains a single point mutation in 2’-O methyltransferase that renders the virus more sensitive to the antiviral effects of interferon (IFN) than wild-type WNV^12^. After intracranial (i.c.) inoculation of WNV-NS5-E218A (WNV), 90% of mice survive acute infection, which is similar to that observed in patients with WNND^13^, allowing the study of neuropathology and behavior in WNV-recovered animals. The viral burden in the brains after WNV infection peaks between 6-8 days post-infection (dpi) (acute phase) and is cleared by 15 dpi. Using this model, we showed that learning and memory deficits that persist after recovery are driven by resident memory CD8 T cell (T_RM_)-IFN-γ-mediated activation of microglia, which eliminate presynaptic termini within the hippocampal trisynaptic circuit^14,15^. Further examination of CD8 T_RM_ in the forebrain revealed that they are maintained in the hippocampus (HPC) via chemokine receptor/ligand CXCR6/CXCL16, and that ∼40% of CD8 T cells also express C-C chemokine receptor type 2 (CCR2)^14^. The functional implications of CCR2 expression by CD8 T cells in the CNS post-flavivirus infection are unknown.

Chemokines and their receptors (CCR2^16^, CCR5^17^, CXCR3^2^, etc.) play an indispensable role in leukocyte trafficking during acute WNV infection. CCR2 is a G-protein-coupled receptor canonically regarded as a chemotactic receptor for monocytes^18^, essential for the effective recruitment of monocytes from the bloodstream to inflammatory sites^19^. During infection with wild-type WNV, CCR2 is critical for monocyte recruitment and viral clearance^16^. However, CCR2 expression by activated CD8 T cells has also been reported in the context of infections with other flaviviruses, including hepatitis C virus and JEV^20,21^, the latter of which infiltrate the CNS and express high levels of programmed cell death protein 1(PD-1). It is unclear whether CCR2^+^ CD8 T cells that persist in the brain after WNV infection promote neuroprotection or neuropathology.

In this study, we characterized the specific cell populations expressing CCR2 via flow cytometry and single-cell RNA sequencing (scRNA-seq) in WNV-infected mice using our model of recovery. Given that both myeloid and lymphoid compartments express CCR2 during recovery, we utilized global CCR2 knockout (KO) mice for adoptive transfer experiments and CD8 T cell-specific CCR2 deletion to delineate the precise function of CCR2^+^ CD8 T cells during acute WNV infection and recovery. While global deletion of CCR2 increased the percentages and cellular levels of CD103, both global and cell-specific depletion of CCR2 consistently led to an increase in the pro-inflammatory functions of CD8 T cells in the hippocampus, including expression of IFN-γ, resulting in worsened recognition memory. Overall, our data indicate a neuroprotective role for CCR2^+^ CD8 T cells that persist as T_RM._

## Results

### CCR2-positive cells persist in the hippocampus in WNV-recovered mice

To define resident and infiltrating mononuclear cell populations that express CCR2 within the forebrain during WNV infection, we utilized Tmem119 ^GFP/+^Ccr2^RFP/+^ mice^22,23^ and i.c. inoculated 10^4^ plaque-forming units (p.f.u.) of WNV. Localization of GFP- and RFP-labeled cells in the HPC during acute WNV infection and after recovery revealed that the majority of RFP^+^ cells enter the dentate gyrus (DG) at 7 dpi (peak infection). At 25 dpi, CCR2^+^RFP^+^ cells were detected within the DG, CA3 and CA1 regions (Fig. 1 A-C, right panel). GFP levels were significantly decreased at 7 dpi across all 3 regions examined (Fig. 1 A-C, left panel), consistent with previous findings demonstrating microglial downregulation of TMEM119 in response to neuroinflammatory conditions^24^. While TMEM119 levels returned to levels comparable to mock-infected animals at 25 dpi (Fig. S1 C), IBA1 levels remained elevated post-infection, indicating a persistent activation of myeloid cells, which include microglia and a small number of persisting macrophages (Fig. 1 A-C, right panel), as previously shown^15^. To better quantitate the kinetics of CCR2^+^ cell subsets during acute WNV encephalitis and after recovery, we examined their dynamics in the forebrains (cortices and hippocampi) of WNV-infected Tmem119 ^GFP/+^Ccr2^RFP/+^ mice via flow cytometry at 7, 15 and 25 dpi (Fig. S1 B). Approximately 30% of CD45^hi^CD11b^+^ cells exhibited single RFP positivity at all timepoints (Fig. 1 E). The observed increase in GFP positivity in CD45^hi^CD11b^+^ and CD45^mid^CD11b^+^ cells was consistent with the sustained activation of microglia, as demonstrated by immunohistochemistry (IHC)^15^. We did observe a small percentage (∼10%) of GFP^+^RFP^+^ CD45^hi^CD11b^+^ cells, which are likely activated microglia, while double-positive CD45^mid^CD11b^+^ cells were negligible at all examined timepoints. The percentage of RFP-positive cells in both CD4 and CD8 T cell populations increased progressively over time with CD8 T cells exhibiting higher RFP expression compared to their CD4 counterparts (Fig. 1 G-H). Overall, these findings established the expression pattern of CCR2 across various mononuclear cell types (CD45^hi^CD11b^+^, CD4 and CD8 T cells) in the forebrain during WNV infection and early recovery.

**Figure 1:**
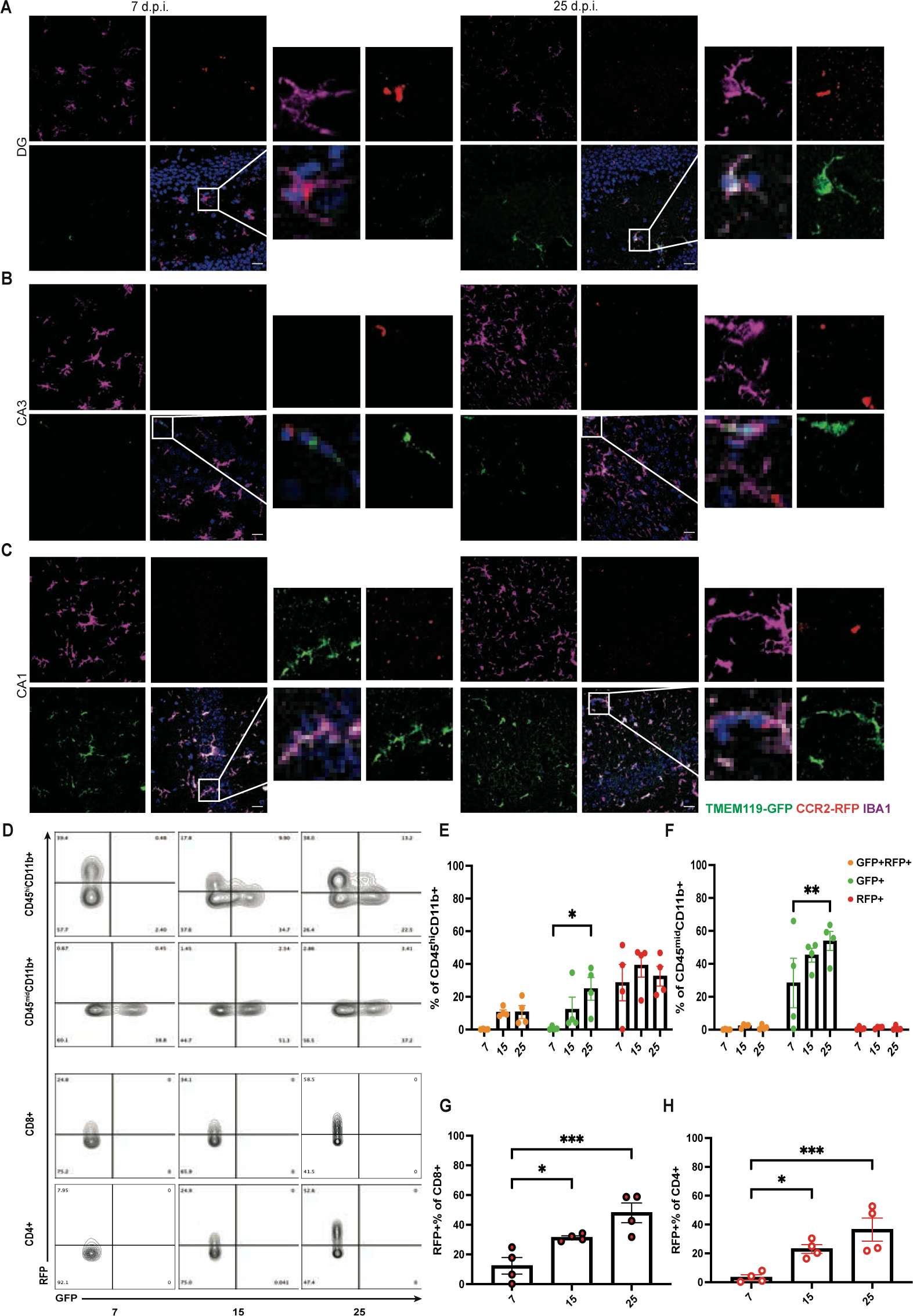
CCR2^+^ cells persist in WNV-recovered HPC. (A-C) Representative immunostaining of DG (A), CA3 (B) and CA1 (C) of hippocampi from Tmem119 ^GFP/+^Ccr2^RFP/+^ reporter mice for GFP (TMEM119), RFP (CCR2) and IBA1 (microglia and macrophage) to identify the source of CCR2 in the hippocampi of WNV-infected mice at 7 (left) and 25 (right) dpi. IBA^+^ area remained elevated throughout viral infection and clearance. Decreased GFP^+^ Sporadic co-localization of GFP and RFP was observed. Images from the boxed area are magnifications of the corresponding images. Scale bars, 20 μm. Data are representative of one independent experiment. (D) Flow cytometry analysis of cells isolated from the forebrains of Tmem119 ^GFP/+^Ccr2^RFP/+^ reporter mice at the indicated dpi (gating strategy shown in Fig. S2 B). GFP^+^ population was not observed in T cells. Data are representative of two independent experiments. (E-H) Quantification of indicated populations (n = 4 mice per group). The percentage of GFP^+^ cells in both CD45^hi^CD11b^+^ and CD45^mid^CD11b^+^ populations gradually increased with viral clearance. Data are representative of one independent experiment. For E-H, data represent the mean ± s.e.m. and were analyzed by two-way ANOVA and corrected for multiple comparisons. *P < 0.05, **P < 0.01, ***P<0.0005.

### Global CCR2 deficiency is associated with up-regulation of IFN-γ and CD103 in CD8 T cells

To examine the functional differences between CCR2^+^ and CCR2^-^ T cells, we further interrogated previously published scRNA-seq data of forebrain immune cells derived from WNV-infected C57BL/6 (wild-type, WT) at 25 dpi^25^. T-distributed stochastic neighbor embedding (t-SNE) plots focused on CD4 and CD8 T cell clusters (Figure 2 A-B) revealed a similar frequency of Ccr2^+^ cells as that observed via flow cytometry of forebrain derived from Tmem119 ^GFP/+^Ccr2^RFP/+^ mice (Fig. 1 G-H), with a higher proportion of CD8 T cells expressing Ccr2 compared to CD4 T cells (Fig. 2 C).

**Figure 2:**
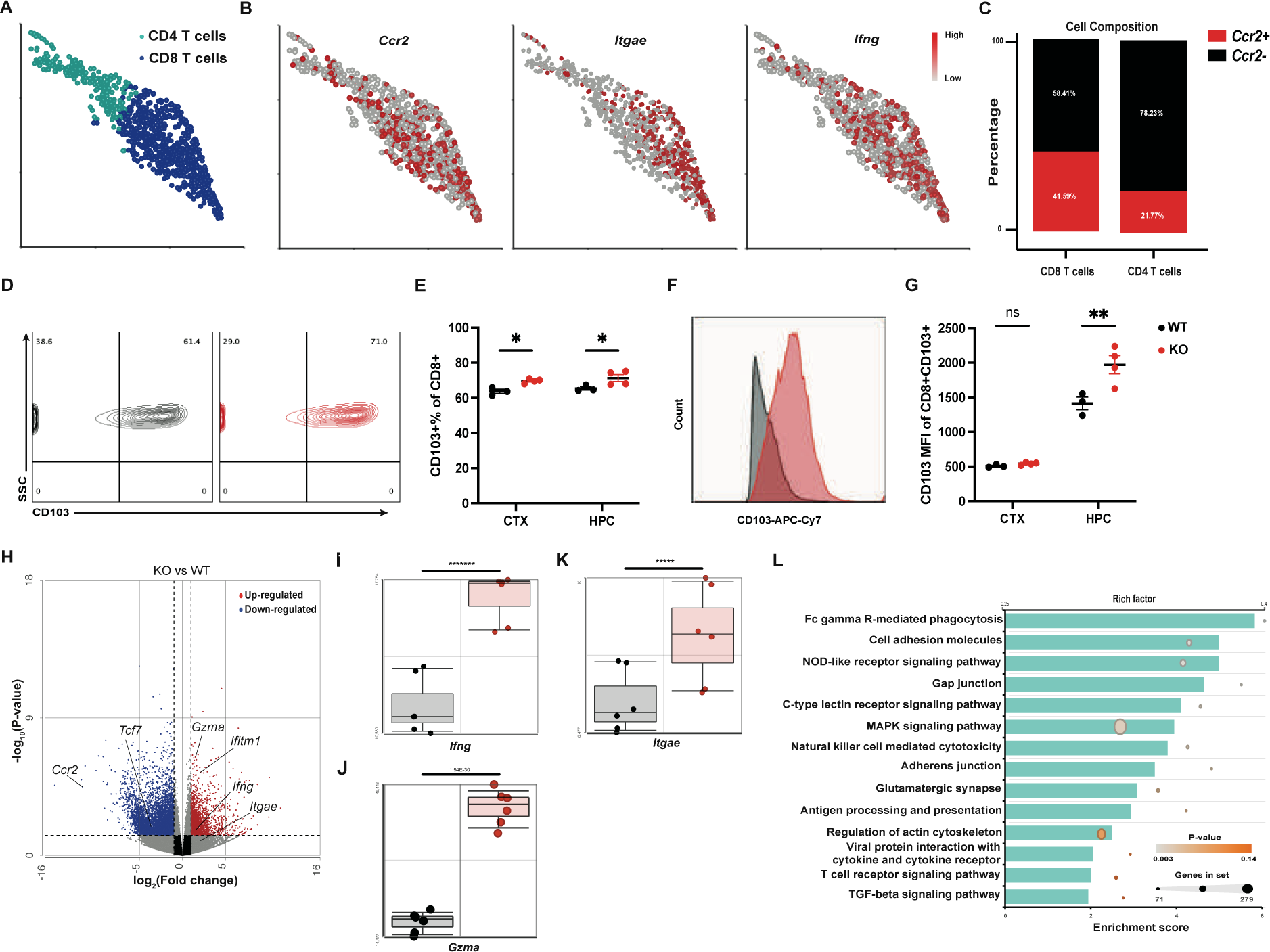
Global CCR2 deficiency contributes to IFN-γ and CD103 up-regulation in CNS CD8 T cells. (A) t-SNE plots focused on CD4 and CD8 T cell clusters from scRNA-seq data of forebrains derived from WNV-infected WT mice at 25 dpi. (B) t-SNE plots with mRNA expression pattern in T cell clusters from WNV-infected WT mice. Ccr2 (left panel), Itgae (middle panel) and Ifng (right panel). Cells are colored by expression value. Values are log-transformed normalized expression counts. (C) Comparison of the proportion of Ccr2^+^ cells between CD4 and CD8 T cells from WT mice at 25 dpi, showing a higher proportion of CD8 T cells expressing Ccr2. (D-G) Flow cytometry analysis demonstrating increased frequency (D-E) and MFI (F-G) of CD103 in CCR2-deficient CD8 T cells compared to WT counterparts. The gating strategy is based on Fig. S1 B followed by a CD8^+^ gate. Data are representative of one independent experiment from 3 repeats. Data represent the mean ± s.e.m. and were analyzed by two-way ANOVA and corrected for multiple comparisons (n = 3 (WT), 4 (KO) mice per group). *P < 0.05, **P < 0.005. ns, not significant. (H) Volcano plot showing DEGs from CD8 T cells isolated from forebrains of WT and KO mice at 30 dpi. Only genes with significant expression level change are shown (adjusted P value < 0.05; log_2_(fold change) < −1 or > 1). Genes with higher expression levels in KO mice are highlighted in red, and genes with lower expression levels in KO mice are highlighted in blue. (I-K) Dot plots showing Ifng, Gzma, Itgae mRNA expression in CD8 T cells from WT and KO animals. *****P < 0.00001, *******P < 0.0000001. (L) Gene Set Enrichment Analysis of DEGs between WT and KO mice using the KEGG database. Only pathways with P value < 0.05 are shown.

To determine whether CCR2 contributes to T cell recruitment and antiviral function within the CNS, we examined immune cell infiltration and viral clearance in WT and Ccr2^RFP/RFP^ (CCR2-deficient, here and after known as KO) mice at 7 and 15 dpi. Global CCR2 deficiency did not affect survival, weight loss or clearance of viral loads in the CNS of WNV-infected animals (Fig. S2 A-D). At 7 dpi, there was a significant reduction in the numbers of CD45^hi^CD11b^+^ cells within the CTX and HPC derived from KO versus WT mice (data not shown). However, no significant differences were observed in the numbers of CD8 T cells (Fig. S2 E). These data indicate that CCR2 is not required for the trafficking and antiviral functions of T cells within the CNS during WNV-NS5-E218A infection.

While CCR2 is not essential for T cell recruitment or virologic control, the increased frequency of CCR2^+^ T_RM_ cells observed at 25 dpi (Fig. 1 G-H) suggests it may play a role in their function during recovery. Canonical cell surface markers for T_RM_ cells include CD69, CD103 and CD44^26^. CCR2-deficient mice exhibited no differences in the frequency of forebrain-derived CD69^+^ and CD44^+^ CD8 T cells, as well as the total number of CD8 T cells, compared to WT mice at 25 dpi (Fig. S2 I-J and H), as well as in the total number of CD8 T cells (Fig. S2 H), suggesting that CCR2 is not essential for T_RM_ maintenance within the virally recovered forebrain. At 25 dpi, CD8 T cell clusters in WT mice express more *Itgae* (CD103) mRNA than CD4 T cell cluster (Figure. 2 B, middle panel). CCR2-deficient CD8 T cells also exhibited increased frequency, cell count and mean fluorescence intensity (MFI) of CD103 (Fig. 2 D-G). Bulk RNA sequencing (RNA-seq) of CD8 T cells isolated from forebrains of WT and KO mice at 30 dpi revealed that genes required for T cell antiviral responses, such as *Gzma* and *Ifng*, were significantly upregulated in the CCR2-deficient versus CCR2^+^ CD8 T cells (Fig. 2 H-J). Increased *Itgae* mRNA expression was also observed, corroborating the results obtained via flow cytometry (Fig. 2 K). Gene Set Enrichment Analysis (GSEA) of differentially expressed genes (DEGs) between WT and KO CD8 T cells derived from the forebrains at 30 dpi using KEGG database (Fig. 2 L) revealed that CCR2 deficiency significantly impacts the T cell receptor (TCR) signaling pathways, leading to an increased percentage of NS4B^+^ T cells (specific for NS4B protein, the immunodominant epitope for WNV in mice^27^) compared with CD8 T cells derived from the forebrain of WNV-infected WT mice (Fig. S2 K). Overall, these data implicate CCR2 signaling in the regulation of CD8 T_RM_ phenotype, including antiviral specificity and IFN-γ expression.

### Extrinsic and intrinsic effects of CCR2 deficiency on CD8 T_RM_ transcriptomic signatures

To study the function of CCR2 specifically on CD8 T cells, we generated *Cd8a*^Cre+^*Ccr2*^fl/fl^ (referred to as Cre^+^ hereafter) animals and Cre^-^ littermate controls^28,29^. *Ccr2*^fl/fl^ mice were constructed to express GFP in Cre^+^ cells where *Ccr2* expression is disrupted. Quantitative reverse transcription polymerase chain reaction (qRT-PCR) analysis of magnetically isolated CD8 T cells and flow-through cells derived from forebrains of WNV-infected Cre^+^ and Cre^-^ mice at 30 dpi confirmed specific *Ccr2* deletion in CD8 T cells (Fig. S3 A). Flow cytometric analysis of these CD8 T cells confirmed their exclusive GFP expression in the forebrain immune cells of WNV-infected Cre^+^ mice (Fig. S3 B). The frequency of GFP^+^ CD8 T cells in WNV-recovered forebrain was approximately 40-50%, similar to that observed in infected *Ccr2*^RFP/RFP^ mice. No significant cell composition change in memory cell subsets was observed upon CCR2 depletion in CD8 T cells (Fig. S3 E-F).

To validate our RNA-seq findings without potentially confounding effects of inactivation of CCR2 on myeloid cells, we performed scRNA-seq analyses on forebrain tissues collected from Cre^+^ and Cre^-^ WNV-infected mice at 30 dpi. We chose an isolation method to enrich myeloid and T cell populations at the expense of most neural cell types (see Methods), which are not the focus of our study. A total of 24,588 cells were collected from 2 biological replicates for each genotype (6 animals for Cre^-^ and 5 animals for Cre^+^) and visualized using a t-SNE plot (Fig. 3 A). We identified 7 major cell types, including B cells, CD4 T cells, CD8 T cells, γδ T cells, microglia, monocytes/macrophages and neural cells (Fig. 3 B and D). Signature genes for each cell cluster were determined by DEGs with the highest fold change over other clusters (Fig. 3 C). DEGs from CD8 T cell clusters were cross-referenced with those from RNA-seq of isolated CD8 T cells (Fig. 3 E). All comparisons were conducted between CCR2-KO samples over CCR2-WT samples. The number of DEGs from RNA-seq analyses of CD8 T cells derived from global KO versus WT mice was greater than that derived from scRNA-seq analysis of Cre^+^ versus Cre^-^ mice, suggesting indirect effects of non-T cells on CD8 T cell transcriptomic signature during global CCR2 deficiency. Examination of overlapping DEGs using GSEA from scRNA-seq revealed that numerous pathways, such as JAK-STAT signaling and TCR signaling, were significantly impacted by CCR2 deletion and consistent with RNA-seq analysis (Fig. 2 L and 3 F). Consistent with previously published scRNA-seq data^25^, CD8 T_RM_ cells are the principal cell type expressing *Ifng* and *Itgae* mRNA post-WNV infection in the forebrain (Fig. S3 G). In Cre^+^ mice, *Ifng* and *Gzma* expressions were increased compared with Cre^-^ mice (Fig. 3 G), confirming the results of RNA-seq analysis in WT and KO mice (Fig. 2 I-J). However, the *Itgae* mRNA level did not change significantly in Cre^+^ animals, in contrast with global KO animals, indicating the extrinsic effects of CCR2 sufficiency by non-T cells on CD103 expression in CD8 T_RM_ cells. IFN-γ-regulating transcription factors *Stat1*^30^, *Stat4*^31^, *Tbx21*^32^ are all increased in Cre^+^ versus Cre^-^ animals (Fig. 3 H), consistent with *Ifng* up-regulation in the context of CCR2-deficiency in both models. Overall, these data revealed transcriptomic signatures in CNS CD8 T cells are regulated both extrinsically and intrinsically by CCR2.

**Figure 3:**
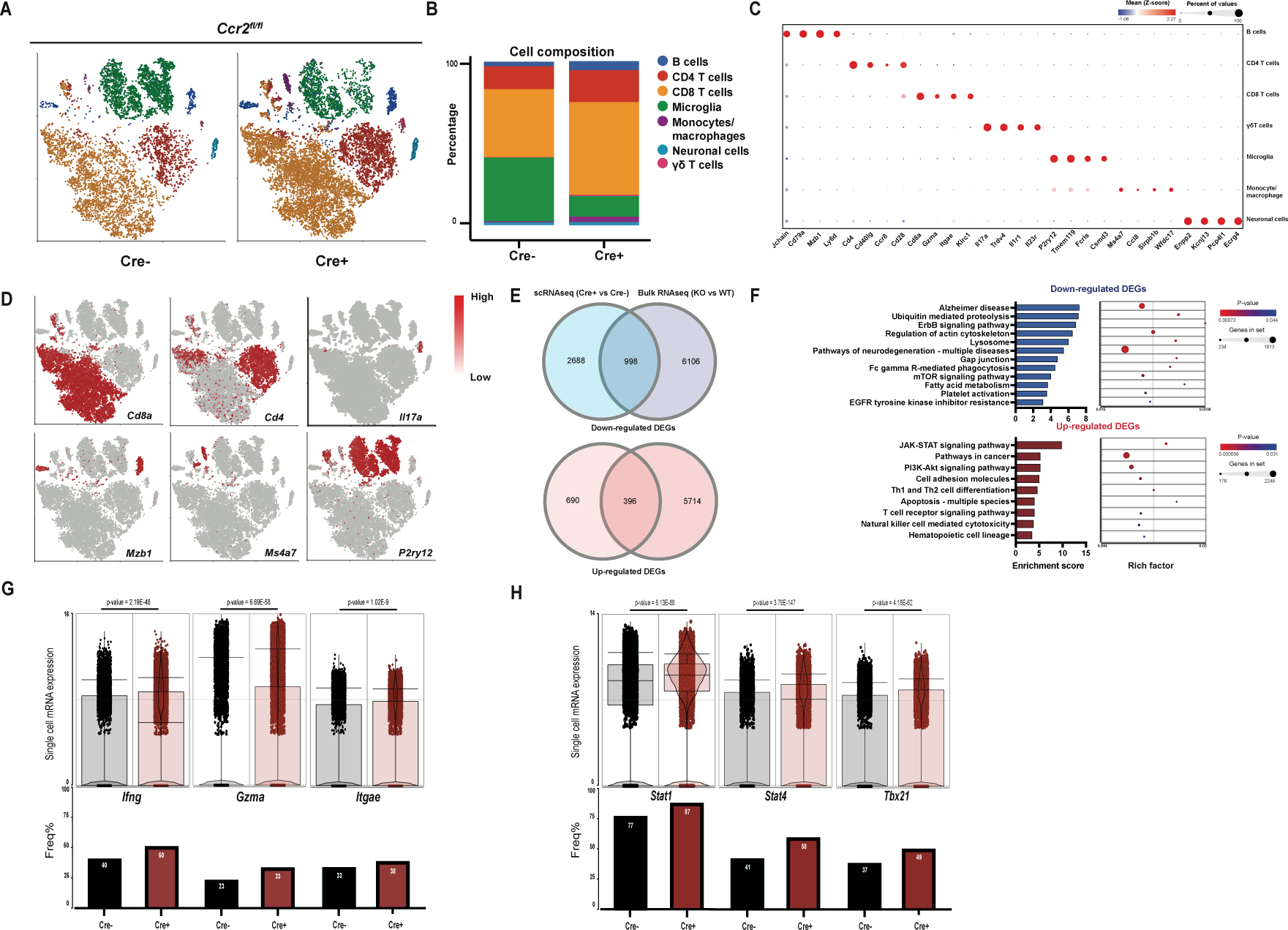
Transcriptomic characterization of CD8-specific CCR2 deficiency post-WNV infection. (A) t-SNE plots visualizing immune cells from the forebrains of WNV-infected Cd8a^Cre+^Ccr2^fl/fl^ (Cre^+^, 5 mice pooled, 2 replicates) and Cd8a^Cre-^Ccr2^fl/fl^ (Cre^-^, 6 mice pooled, 2 replicates) mice at 30 dpi. Clusters are colored by cell-type identity as shown in the legend next to Fig. 3 B. The plots show the distribution of seven major cell types: B cells, CD4 T cells, CD8 T cells, γδ T cells, microglia, monocytes/macrophages, and neurons. Cell types were annotated using a combination of DEGs and previously published data as reference. (B) Proportion of every identified cell type in each genotype at 30 dpi. (C) Heatmap of signature genes for each identified cell cluster, determined by DEGs with the highest fold-change. (D) Expression pattern of canonical marker genes for each cell type (neurons excluded), including Cd8a (CD8 T cells), Cd4 (CD4 T cells), Il17a (γδ T cells), Mzb1 (B cells), Ms4a7 (monocytes/macrophages) and P2ry12 (microglia). Cells are colored by expression value. Values are log-transformed normalized expression counts. (E) Venn diagram of up- and down-regulated DEGs from the scRNA-seq and bulk RNA-seq of WT and KO mice. DEGs from scRNA were generated by ANOVA differential analysis on Cre^+^ over Cre^-^ animals. DEGs from bulk RNA-seq were generated by DEseq2 differential analysis on KO over WT animals. A total of 998 genes were down-regulated, and 396 genes were up-regulated in both sequencing experiments. (F) Biological pathways from GSEA based on overlapped DEGs from Fig. 3E, revealing significant impacts on pathways such as JAK-STAT signaling and multiple diseases of neurodegeneration due to CCR2 deficiency on CD8 T cells. (G-H) Upper panel: violin plots of genes of interest from scRNA-seq. Lower panel: frequency of cells expressing RNA of genes of interest from scRNA-seq.

### CCR2 limits IFN-γ expression by CD8 T_RM_ cells within the hippocampus

While scRNA-seq analyses of Cre^+^ versus Cre^-^ animals revealed significant transcriptomics changes and enabled us to identify distinct cell types and subpopulations within the CNS, there are technical limitations to scRNA-seq, such as low transcript capture efficiency and sequencing coverage^33^. To validate the DEGs from scRNA-seq with higher confidence, we performed bulk RNA-seq on forebrain tissues isolated from WNV-infected Cre^+^ and Cre^-^ littermates at 30 dpi. (Fig. 4A)

**Figure 4:**
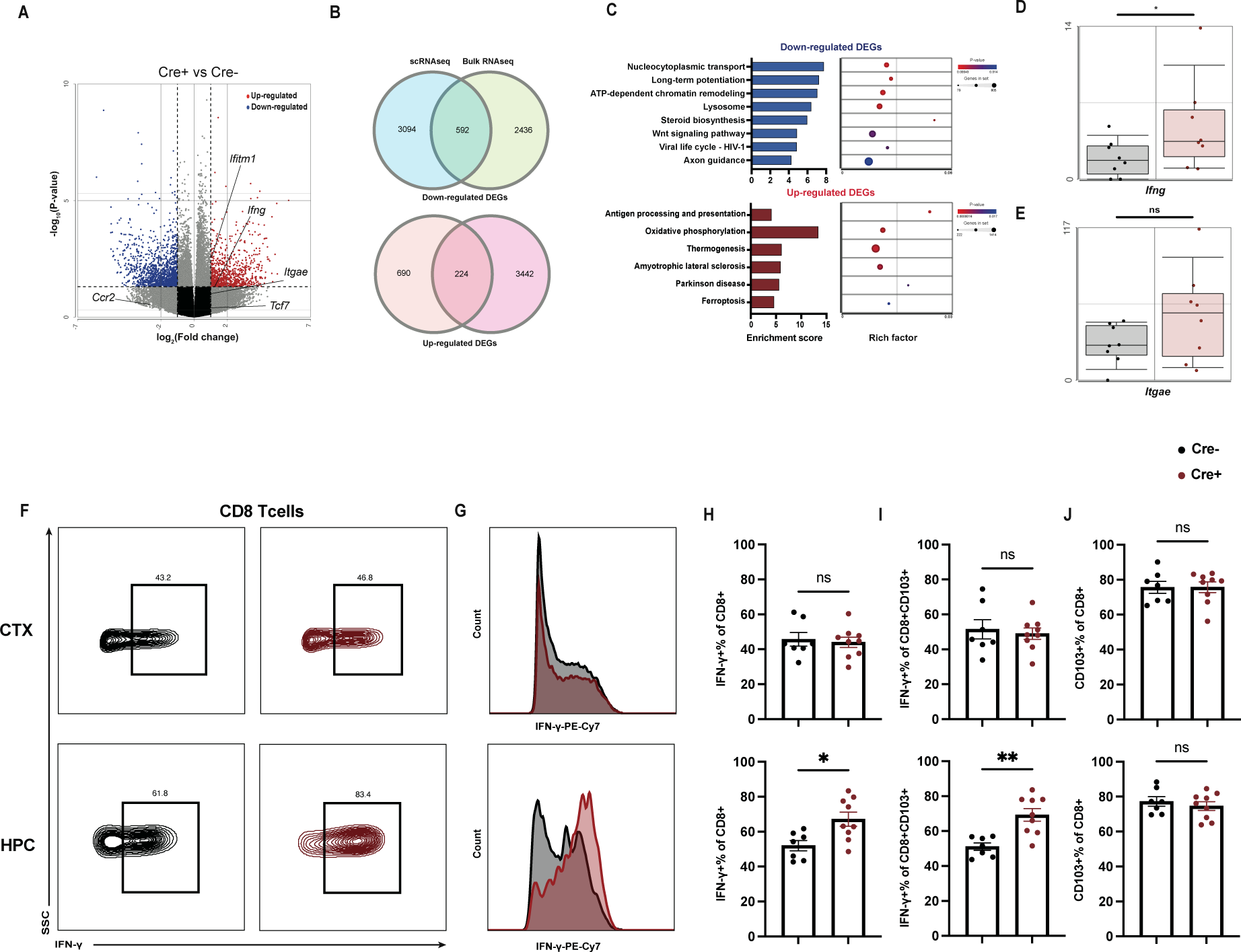
IFN-γ production is elevated by CCR2 deficiency in hippocampal CD8 T cells. (A) Volcano plot showing DEGs from forebrains of WNV-infected Cre^+^ and Cre^-^ mice at 30 dpi. Only genes with significant expression level change are shown (adjusted P value < 0.05; log_2_(fold change) < −1 or > 1). Genes with higher expression levels in Cre^+^ mice are highlighted in red, and genes with lower expression levels in Cre^+^ mice are highlighted in blue. (B) Venn diagram of up- and down-regulated DEGs from the scRNA-seq and bulk RNA-seq of Cre^+^ and Cre^-^ mice. DEGs from bulk RNA-seq were generated by DEseq2 differential analysis on Cre^+^ over Cre^-^ animals. A total of 592 genes were down-regulated, and 224 genes were up-regulated in both sequencing experiments. (C) Biological pathways from GSEA based on overlapped DEGs from Fig. 4 B, revealing significant impacts on pathways such as lysosome and antigen processing and presentation due to CCR2 deficiency on CD8 T cells. (D-E) mRNA expression levels of Ifng and Itgae from bulk RNA-seq. Ifng showed consistent up-regulation across both bulk RNA-seq and scRNA-seq, while Itgae mRNA levels were inconclusive. (F-H) Flow cytometry analysis demonstrating increased frequency (F and H) and MFI (G) of IFN-γ in CD8 T cells from HPC of Cre^+^ mice compared to Cre^-^ littermates at 30 dpi. CD8 T cells from CTX were not affected by CCR2 deficiency. The gating strategy is based on Fig. S1 B followed by a CD8^+^ gate. (I) Quantification of the percentage of CD103^+^ CD8 T cells that are IFN-γ^+^. (J) Quantification of the percentage of CD8 T cells that are CD103^+^. For F-J, data were pooled from two independent experiments (n = 7 (Cre^-^), 9 (Cre^+^) mice per group). Data represent the mean ± s.e.m. and were analyzed by unpaired t-tests. *P < 0.05, **P < 0.005. ns, not significant.

DEGs identified via bulk RNA-seq of forebrains from WNV-recovered Cre^+^ and Cre^-^ animals were cross-referenced with those from scRNA-seq analysis (Fig. 4 B). A total of 592 genes were down-regulated and 224 genes were up-regulated in both sequencing analyses. GSEA analysis revealed significant alterations in the genes within the lysosome, neurodegeneration, T cell receptor signaling, and antigen processing and presentation pathways, which were significantly affected by the manipulation of CCR2 (Fig. S4 C). In addition, the expression of *Ifng* mRNA in the forebrain derived from Cre^+^ mice was significantly up-regulated compared with Cre^-^ animals, consistent with bulk RNA-seq performed on CD8 T cells derived from WT and KO mice and scRNA-seq of forebrain from Cre^+^ and Cre^-^ mice (Fig. 4 A, D). Notably, the *Itgae* mRNA levels did not differ in both bulk RNA-seq and scRNA-seq studies performed on Cre^+^ and Cre^-^ animals, which further confirms that CD103 up-regulation observed in KO mice is likely to be extrinsically induced (Fig. 4 E). Finally, flow cytometric analyses of forebrain tissues derived from WNV-recovered Cre^+^ and Cre^-^ mice at 30 dpi revealed significantly higher percentages of IFN-γ^+^CD103^+^ CD8 T_RM_ cells in the HPC, but not the CTX, of Cre^+^ mice, and higher levels of IFN-γ expression per cell compared to that of Cre^-^ mice (Fig. 4 F-H). Furthermore, the same observation could be made in CD103^+^ CD8 T cells (Fig. 4 I) but not in CD103^-^ CD8 T cells (Fig. S4 E), which suggests that CD103^+^ CD8 T cells in Cre^+^ mice were specifically contributing to the elevated IFN-γ levels in the HPC. Of note, the overall percentages of CD103^+^ CD8 T cells were not significantly different between the genotypes in CTX and HPC (Fig. 4 J), corroborating findings in both bulk RNA-seq and scRNA-seq analyses of WNV-recovered forebrains of Cre^+^ and Cre^-^ animals.

### CCR2 intrinsically regulates IFN-γ expression in virus-specific CD8 T cells

To validate the intrinsic effect of CCR2 on CD8 T cell function *in vivo*, we performed adoptive transfer experiments. TCR-transgenic mice with specificity for the WNV immunodominant epitope in NS4B (here and after known as WNV-I mice^34^) were crossed with CCR2 KO mice to generate CCR2-deficient WNV-I mice. Leukocytes of WNV-I mice express CD45.1, allowing them to be distinguished from those of WT recipient animals, which express CD45.2 (Fig. S5 A). Naïve CD8 T cells isolated from the spleens of WT WNV-I or CCR2-deficient WNV-I mice were adoptively transferred into WT recipient mice 24 hours before WNV infection and forebrains of recipient mice were harvested for flow cytometric analyses at 30 dpi. MHC class I tetramer staining confirmed that nearly 100% of the transferred CD8 T cells were NS4B^+^ (Fig. S5 B and D), as expected. Forebrain retention of CD8 T cells was not affected by CCR2 deficiency, confirming that CCR2 is not responsible for T_RM_ maintenance in the CNS post-WNV infection (Fig. S5 C and E). Notably, both IFN-γ positivity and MFI levels were higher in CCR2-deficient T cells compared to those expressing CCR2 (Fig. 5 C and F). Additionally, almost all the transferred WNV-I CD8 T cells expressed CD103, which is much higher than that observed in previous experiments, and consistent with the finding that WNV-recovered KO mice exhibit increased CD103 expression and NS4B compared with their WT counterparts (Fig. 2 E and S2 K). Overall, these results confirmed that CCR2 intrinsically regulates IFN-γ production by CNS CD8 T cells post-WNV infection.

**Figure 5:**
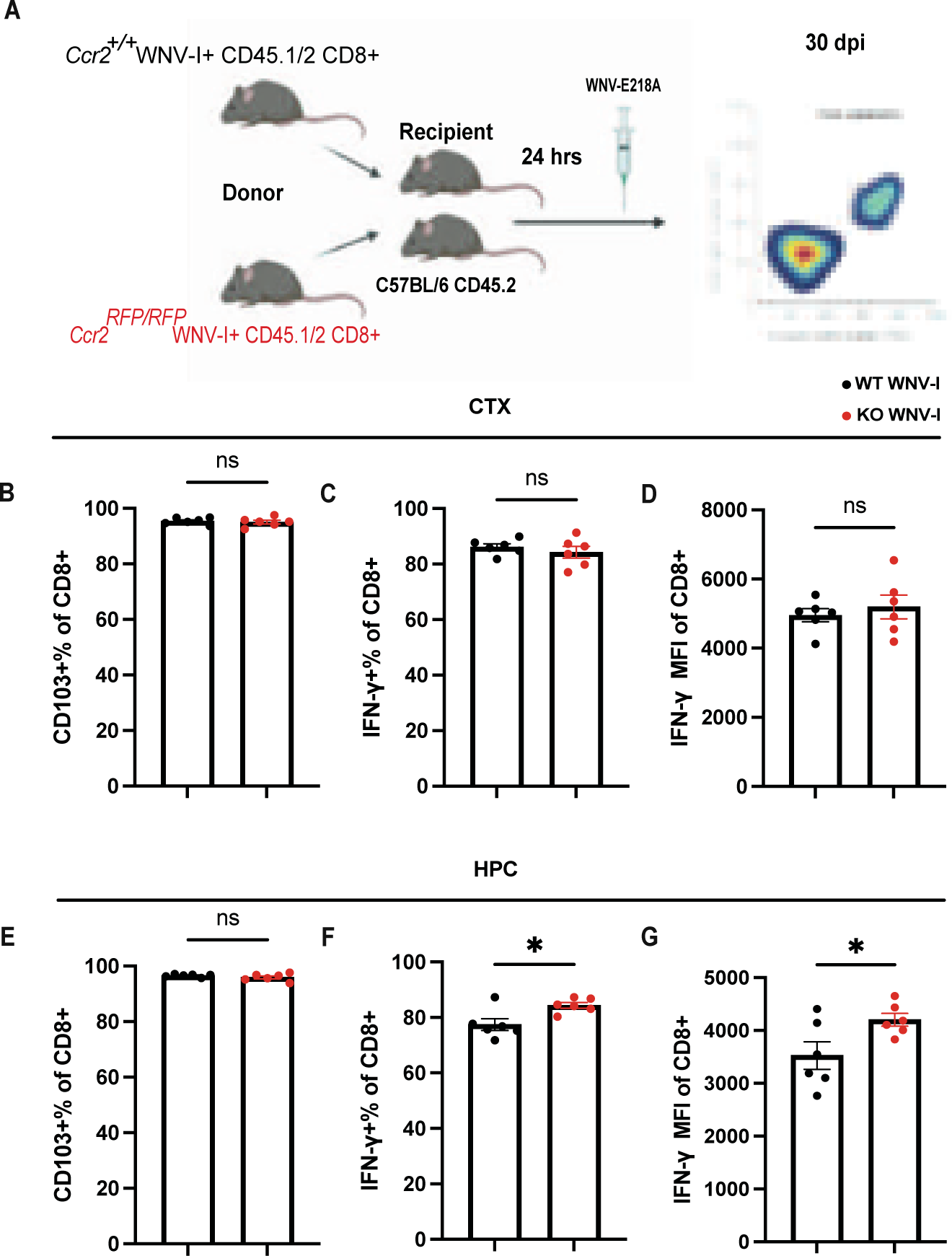
CCR2 intrinsically regulates IFN-γ expression of hippocampal CD8 T cells. (A) Schematic showing the experimental design of CD8 T cell adoptive transfer from WT and KO CD45.1/2 WNV-I mice into WT (CD45.2) mice. 5, 000 CD8 T cells were retro-orbitally injected 24 hours before WNV infection. CTX and HPC of the recipient mice were harvested at 30 dpi for flow cytometry. (B and E) Quantification of the percentage of transferred CD8 T cells that are CD103^+^. T_RM_ from the transgenic mice exhibited much higher CD103 expression than those from WT mice. (C and F) Quantification of the percentage of transferred CD8 T cells that are IFN-γ^+^. (D and G) MFI of IFN-γ of transferred CD8 T cells showing higher IFN-γ expression in CD8 T cells from the HPC. Data are representative of one independent experiment from 3 repeats. Data represent the mean ± s.e.m. and were analyzed by unpaired t-tests (n = 6 (WT WNV-I), 6 (KO WNV-I) mice per group). *P < 0.05. ns, not significant.

### CCR2-deficient CD8 T cells worsen novel object recognition post-WNV infection

We previously showed that T cell-derived IFN-γ contributes to hippocampal-based memory impairments during recovery from WNV infection^14^. Given that CCR2 inactivation in CD8 T cells leads to increased IFN-γ expression within the HPC, we hypothesized that CCR2 deficiency-induced IFN-γ may negatively affect cognitive recovery. To test this, WNV-recovered Cre^+^ and Cre^-^ littermates were subjected to Novel Object Recognition (NOR) testing, a two-day test which investigates various aspects of learning and memory in mice (Fig. 6 A). During the habituation phase, no difference in total mobile time was observed, indicating that locomotor activity was unaffected by CD8-specific CCR2 deletion (Fig. 6 B). On the training day, both groups spent similar amounts of time with two identical objects, showing no preference for either (Fig. 6 C). The number of investigations was also comparable between the groups, further confirming no preference between the identical objects. During the testing phase, Cre^-^ mice exhibited a significantly greater preference for the novel object, as shown by the absolute discrimination measure (Fig. 6 E). Cre^-^ mice also spent significantly more time interacting with the novel object, whereas Cre^+^ mice displayed no significant difference in exploration between the novel and familiar objects (Fig. 6 F). The number of investigations of the novel object by Cre^-^ mice was significantly higher than that of Cre^+^ mice (Fig. 6 G). Overall, these findings suggest that CCR2 signaling in CD8 T cells preserves recognition memory following WNV infection.

**Figure 6:**
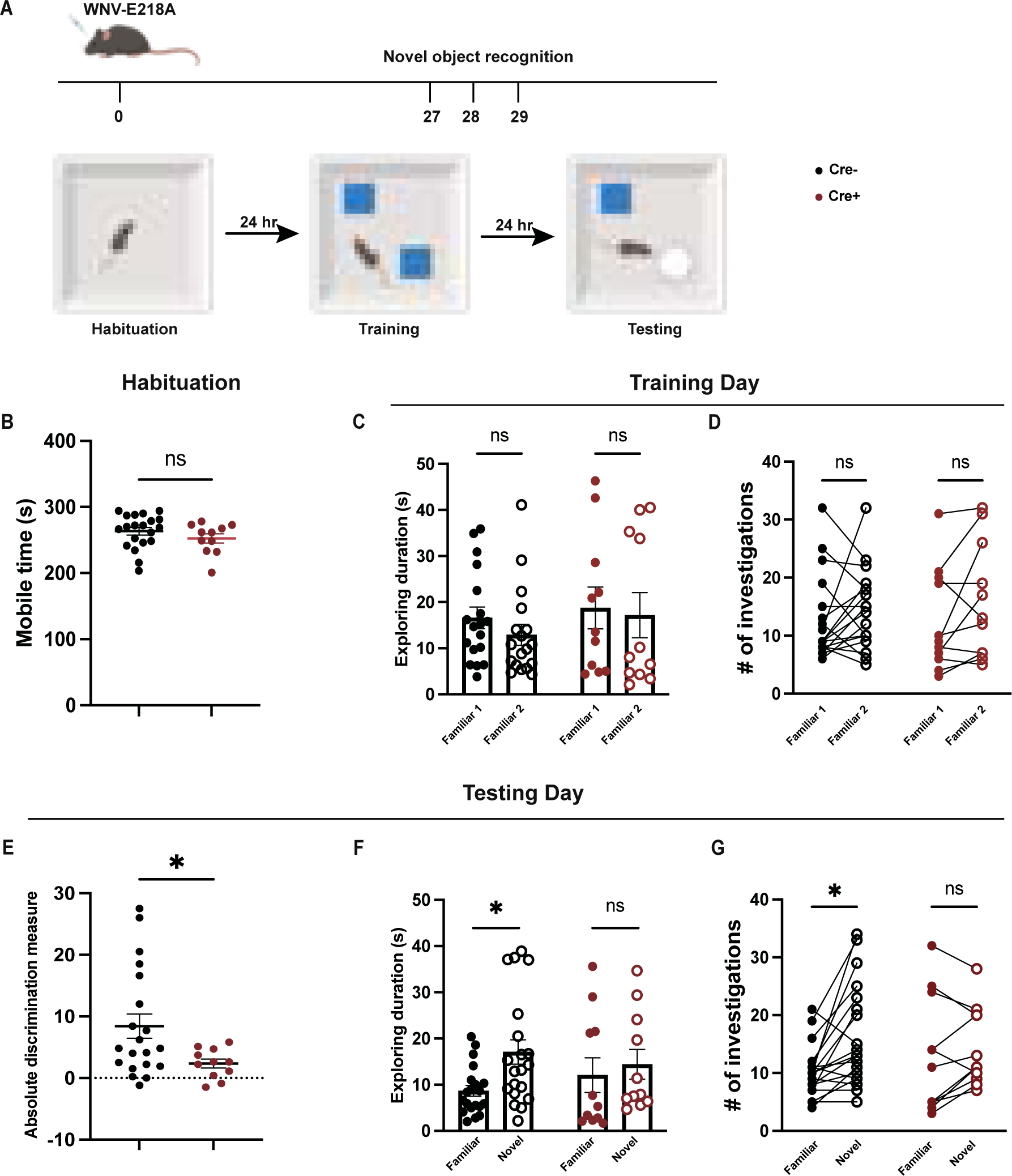
Impact of CCR2 deficiency on recognition memory during WNV recovery. (A) Schematic showing the experimental design of the novel object recognition test to assess non-spatial recognition memory in WNV-infected Cre^+^ and Cre^-^ littermates. (B) The total mobile time during the habituation phase showed no significant difference between Cre^+^ and Cre^-^ mice, indicating comparable locomotor activity. (C) Time spent and (D) Number of interactions with two identical objects during the training phase. (E) Absolute discrimination measure, showing that Cre^-^ mice had a significantly greater preference for the novel object compared to Cre^+^ mice. (F) Time spent and (G) Number of interactions with novel and familiar objects during the testing phase. For B and E, data were analyzed by unpaired t-tests. For C-D and F-G, data were analyzed by two-way ANOVA and corrected for multiple comparisons. (n = 20 (Cre^-^), 11 (Cre^+^) mice per group). Data were pooled from 3 independent experiments. *P < 0.05. ns, not significant.

## Discussion

Chemokines and their receptors are an integral part of antiviral responses in the CNS^2,16,17^. Although many studies have demonstrated that T cells express CCR2 in the context of neuroinflammation, its functional implications for T cells remain unclear. Here, we investigated the role of CCR2 expression on CD8 T cells in the CNS post-WNV infection, exploring how this chemokine receptor influences T_RM_ maintenance, immune responses and cognitive outcomes during recovery from viral encephalitis. Using a dual-reporter mouse model to delineate CCR2 expression by cell-specific markers, we found that CCR2 is expressed by CD45^hi^CD11b^+^, CD4 and CD8 T cells in the forebrain during acute infection and early recovery from WNV. Comparisons between sequencing experiments of forebrain immune cells derived from global CCR2 KO versus CD8-specific CCR2-deficient mice were utilized to define intrinsic versus extrinsic effects of CCR2 on CD8 T cell functions in the forebrains post-WNV infection. We found that CCR2 is not required for the trafficking or antiviral functions of CD8 T cells during acute infection, or the maintenance of CD8 T_RM_ during recovery, within the WNV-NS5-E218A-infected forebrain. The lack of effect of CCR2 expression on virologic control contrasts with that observed in studies utilizing CCR2 KO mice during infections with WT WNV^16^. This is likely due to the enhanced effects of type I interferon-mediated innate immune responses to effectively clear WNV-NS5-E218A.

Despite the lack of effect of CCR2 on T cell recruitment and virologic control, we observed increased IFN-γ and CD103 expression levels in CD8 T cells derived from global KO compared to WT CD8 T cells during recovery, suggesting this chemokine receptor limits T_RM_ differentiation and pro-inflammatory function. This is consistent with a study examining influenza A infection in which lung T_RM_ showed enhanced expression of CD103 in CCR2-deficient CD8 T cells^35^. CD8-specific CCR2-deficient mice exhibited only increased IFN-γ production, suggesting that CD103 up-regulation may be extrinsically induced by CCR2 signaling in the broader immune environment. Given that CCR2 is expressed on both lymphoid and myeloid cells, global depletion of CCR2 could alter cytokine productions from both CD4 T cells and macrophages, which in turn led to CD103 up-regulation on CD8 T cells. Adoptive transfer experiments with CD8 T cells further confirmed that CCR2 intrinsically regulates IFN-γ production by CD8 T cells in the WNV-recovered forebrain. Increased IFN-γ levels induced by CCR2 depletion on CD8 T cells worsened the recognition memory deficits in WNV-recovered mice.

Our study also reveals that CCR2-positive cells persist in the CNS throughout the recovery phase post-WNV infection, with a significant presence in both CD4 and CD8 T cell populations. The persistence of these cells indicates that CCR2 may be involved in the immune response in the CNS post-WNV infection. By using global CCR2 KO mice and CD8 T cell-specific CCR2 deletion models, we observed that while CCR2 is not essential for T cell recruitment or antiviral functions, it significantly influences the phenotype and function of CD8 T cells within the CNS. There have been conflicting reports on whether microglia express CCR2^36–38^ upon inflammation. We found that there were only negligible Tmem119-GFP^+^CCR2-RFP^+^ CD45^hi^CD11b^+^ cells present in the CNS post-viral clearance.

Through scRNA-seq and bulk RNA-seq analyses, we identified significant transcriptomic changes in CCR2-deficient CD8 T cells, including upregulation of genes involved in T cell antiviral responses, such as *Gzma* and *Ifng*. Gene Set Enrichment Analysis (GSEA) revealed that CCR2 deficiency impacts key signaling pathways, including JAK-STAT and T cell receptor (TCR) signaling. These findings underscore the importance of CCR2 in modulating T cell activation and function within the CNS during recovery from WNV infection.

Our behavioral studies using the NOR test revealed that CD8 T cell-specific deletion of *Ccr2* (Cre^+^ mice) significantly worsens recognition memory post-WNV infection. Previous studies have shown that IFN-γ KO animals perform better in novel object recognition (NOR) tests^39^. Cre^+^ mice showed impaired performance in the NOR test compared to their Cre^-^ littermates. This cognitive impairment was associated with increased IFN-γ production by hippocampal CD8 T cells, consistent with prior studies indicating that IFN-γ levels contribute to the memory deficits.

In summary, our study highlights the critical role of CCR2 in regulating CD8 T cell function and neuroinflammation in the CNS following WNV infection. CCR2-deficient CD8 T cells in the CNS exhibited elevated IFN-γ production, which contributed to cognitive impairments during the recovery from viral encephalitis. These findings offer important insights into the mechanisms of T cell-mediated neuroinflammation. Our results also suggest the need for caution when considering the use of CCR2 antagonists for other diseases, as they could inadvertently cause neurological side effects by disrupting CCR2’s essential regulatory function in the CNS.

## Materials and methods

### Mouse strains

All experiments followed guidelines approved by the Washington University School of Medicine Animals Safety Committee (protocol number 21-0019). Eight to 12-week-old male and female mice were used for all experiments. C57BL/6J, Ccr2^RFP/RFP^ (017586), Tmem119^GFP/GFP^ (031823) and Cd8a^cre+^ (008766) mice were purchased from Jackson Laboratories. Ccr2^fl/fl^ mice were obtained from Dr. Manolis Pasparakis (University of Cologne) and Dr. Yu-qing Cao (Washington University). WNV-I transgenic mice were obtained from Dr. Michael Diamond (Washington University).

### Mouse model of WNV infection

WNV-NS5-E218A strain used in this study contains a single point mutation in the 2′-O-methyltransferase and was obtained from Michael Diamond (Washington University)^12^. Mice were deeply anesthetized with a cocktail of ketamine/xylazine/acepromazine and intracranially administered 10^4^ plaque-forming units (p.f.u.) of WNV-NS5-E218A into the brain’s third ventricle using a guided 29-gauge needle. The virus was diluted in 10 μL of 0.5% fetal bovine serum in Hank’s balanced salt solution (HBBS, Gibco). The mock-infected mice were administered with the virus diluent alone.

### Immunohistochemistry

Mice were anesthetized and sacrificed by cardiac perfusion with ice-cold dPBS (Gibco), followed by ice-cold 4% paraformaldehyde (PFA). Brain tissues were dissected and immersed in 4% PFA overnight, followed by immersion in 30% sucrose (freshly changed every 24h, 3 times). Prior to tissue sectioning (10 μm), samples were frozen in OCT compound (Fischer Scientific, cat. no. 23-730-571). Tissue sections were washed with PBS and permeabilized with 0.1% Triton X-100 (Sigma-Aldrich) and blocked with 5% normal goat serum (Sigma-Aldrich) at room temperature for 1h. Slides were then incubated in primary antibody overnight at 4°C. Following 3 washes in PBS, slides were then incubated in secondary antibodies at room temperature for 1 h, and nuclei were counterstained with 4,6-diamidino-2-phenylindole (DAPI; Invitrogen). Coverslips were applied with ProLong Gold Antifade Mountant (Invitrogen). Immunofluorescent images were acquired using a Zeiss LSM 880 confocal laser scanning microscope and processed using software from Zeiss. Immunofluorescent signals were quantified using the software ImageJ.

### Antibodies for immunohistochemistry

The following primary antibodies were used for IHC analyses: IBA1 (1:200; Synaptic Systems, cat. no. 234009), synaptophysin (1:250; Synaptic Systems, cat. no. 101004) and Homer1 (1:250; Synaptic Systems, cat. No. 160003). Secondary antibodies conjugated to Alexa-555 (Invitrogen, cat. no. A21437 and A21435) or Alexa-647 (Invitrogen, cat. no. A32933) were used at a 1:200 dilution.

### Flow cytometry

Cells were isolated from murine hippocampi and cortices at indicated dpi and stained with fluorescence-conjugated antibodies (described below). In brief, mice were anesthetized and sacrificed by cardiac perfusion with ice-cold dPBS (Gibco). Brains were aseptically dissected, minced and enzymatically digested in HBSS containing 0.05% collagenase D (Sigma), 0.1 μg/mL TLCK trypsin inhibitor (Sigma), 10 μg/mL DNase I (Sigma), and 10 mM HEPES (Gibco) at 37°C for 1h with shaking. Tissues were then pushed through a 70 μm strainer and centrifuged at 500× g for 10 min. Brain cell pellets were resuspended in 37% isotonic Percoll (Cytiva, cat. no. 17089101) and centrifuged at 1200× g for 30 min to remove myelin debris. Cells were washed in PBS. For cytokine restimulation, isolated cells were then treated with Cell Activation Cocktail (BioLegend, cat. no. 423304) to stimulate cytokine expression for 4 h at 37 °C, 5% CO_2_. Prior to immunostaining, all cells were blocked with TruStain FcX antibody (Biolegend, cat. no. 101320) for 5 min. Cells were stained with LIVE/DEAD Fixable Blue Dead Cell Stain Kit at 1:1000 dilution following dissolution in 50 μL DMSO (Invitrogen, cat. no. L34962) and extracellular antibodies for 30 min at room temperature, then washed twice with PBS, fixed with 2% PFA for 10 min at room temperature. Intracellular markers were detected following extracellular staining. Cells were permeabilized with eBioscience Foxp3/Transcription Factor Staining Buffer Set (Invitrogen, cat. no. 00-5523-00) and stained with intracellular antibodies for 30 min at room temperature, then washed twice with permeabilization buffer. Data were collected with a BD LSR Fortessa X-20 flow cytometer and analyzed with FlowJo software.

### Antibodies for flow cytometry

The following antibodies were used for flow cytometry: CD45 (BUV737, BD, cat. no. 748371), CD8 (Brilliant Violet 711, BioLegend, cat. no. 100748), CD103 (APC/Cyanine7, BioLegend, cat. no. 121432), IFN-γ (PE/Cyanine7, BioLegend, cat. no. 505826), CD69 (BV421, BD, cat. no. 562920), CD11b (Brilliant Violet 605, BioLegend, cat. no. 101257), CD44 (APC, BioLegend, cat. no. 103011), CD45.1(APC, BioLegend, cat. no. 110713) and CD45.2 (BUV737, BD, cat. no. 612779). WNV-specific CD8 T cells were identified with H-2D(b) WNV NS4b 2488-2496 SSVWNATTA Brilliant Violet 421-labeled tetramer obtained from the NIH Tetramer Facility.

### Measurement of viral burden

Mice were infected with WNV and euthanized at specific dpi, as indicated. For tissue collection, mice were deeply anesthetized and underwent cardiac perfusion. All tissues collected were weighed and then homogenized in 500 μL of PBS and stored at -80°C until virus titration. Viral titers were quantified as described previously^14^.

### Single-cell RNA sequencing

ScRNA-seq was performed on Cre^-^ (6 mice, 2 replicates) and Cre^+^ (5 mice, 2 replicates) mice. Brain cell pellets were obtained as described in the Flow cytometry section following Percoll gradient centrifugation. Dead cells were removed using the Dead Cell Removal Kit (Miltenyi Biotec, cat. no. 130-090-101). Live cells were diluted to 1000 cells/µL with ice-cold 0.04% BSA in PBS and sent to Genome Access Technology Center at the McDonnell Genome Institute for library preparation using 10x Genomics Chromium Single Cell 5’ Reagent Kits and sequencing. Data was analyzed in Partek Flow software v12.0.2 suite with the standard pipeline for scRNA-seq analyses to conduct QA/QC filtering, align and normalize reads, identify cell clusters, generate lists of DEGs and conduct GSEA.

### Bulk RNA sequencing

For WT and CCR2 KO mice, CD8 T cells from the CNS were enriched from brain cell pellets post Percoll gradient centrifugation using MojoSort Mouse CD8 T Cell Isolation Kit (BioLegend, cat. no. 480035). For Cre^-^ and Cre^+^ mice, forebrain tissues were collected following cardiac perfusion. RNA was isolated from these cells/tissues with RNeasy kit (QIAGEN, cat. no. 74104 and 74004). Samples underwent polyA selection and prepared according to the library kit manufacturer’s protocol, indexed, pooled, and sequenced on an Illumina NovaSeq X Plus system. Data was analyzed in Partek Flow software v12.0.2 suite with the standard pipeline for bulk RNA-seq analyses to conduct QA/QC filtering, align reads, normalize transcript counts, generate lists of DEGs and conduct GSEA.

### qRT-PCR

For Cre^-^ and Cre^+^ mice, CD8 T cells from the CNS were enriched from brain cell pellets post Percoll gradient centrifugation using MojoSort Mouse CD8 T Cell Isolation Kit. After RNA concentration was measured using Nanodrop 2000 instrument (ThermoFisher), MultiScribe Reverse Transcriptase (Invitrogen, cat. no. 4311235) was used to generate cDNA. qRT-PCR was carried out using Power SYBR PCR Master Mix (Applied Biosystems, cat. no. 4367659) with target-specific primers on a Bio-Rad CFX384 instrument according to manufacturer recommendations. The forward and reverse primers were as follows: Ccr2, 5′-ATCCACGGCATACTATCAACATC-3′ and 5′-CAAGGCTCACCATCATCGTAG-3′.

### CD8 T cell adoptive transfer experiments

Spleens harvested using the aseptic technique from naïve WT or CCR2 KO WNV-I mice were placed in sterile, ice-cold PBS and then mushed through a 70 μm strainer. CD8 T cells were isolated using the MojoSort Mouse CD8 T Cell Isolation Kit (BioLegend, cat. no. 480035). T cells were washed with fresh PBS 3 times through centrifugation. 5,000 CD8 T cells resuspended in 150 μL PBS were retro-orbitally injected into a WT recipient.

### Novel object recognition test

During the habituation phase, mice were placed in an empty open arena for 5 min. 24 h later, mice were placed into the same arena with 2 identical objects fixed to the floor in the opposite corner of the box. During the training day, mice were allowed to explore for 5 min. 24 h later, one familiar object was replaced by a novel object (different in both color and shape). During the testing day, mice were allowed to explore for 5 min. All trials were video-recorded using a camera and analyzed with ANY-maze software. (Absolute discrimination measure = novel object exploration time − familiar object exploration time)

## Acknowledgments

We thank Dr. M. Diamond (Washington University) for the gifting of WNV-I mice, and Drs. Y. Cao (Washington University) and Dr. M. Pasparakis (University of Cologne) for sharing *CCr2^fl/fl^* mice for our study. We also thank W. Beatty at the Molecular Microbiology Imaging facility at Washington University School of Medicine. Special thanks to Dr. A. Kravitz and J. Ma for their assistance in 3D printing. This study was funded by R35NS122310 (to R.S.K.).

---

**Figure S1:**
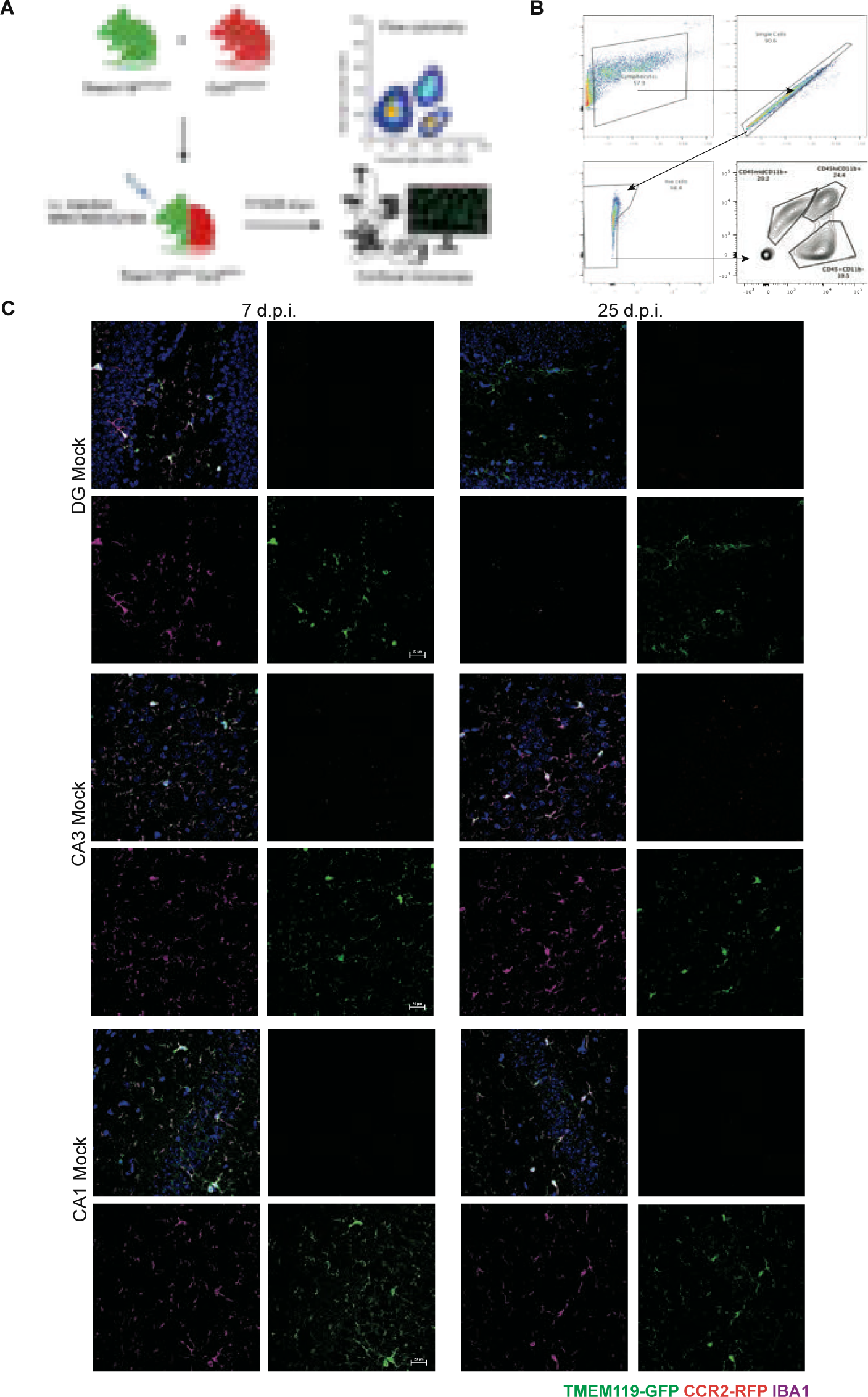
The Tmem119 ^GFP/+^Ccr2^RFP/+^ reporter mouse is a great tool for visualizing and quantifying CCR2 expression. (A) Experimental design for reporter mouse breeding, infection method and identifying immune cell subsets for CCR2 expression. Mice were infected intracranially with 1×10^4^ p.f.u. WNV-NS5-E218A and harvested at indicated timepoints. (B) Gating strategy for flow cytometry. Cells were gated via CD45 and CD11b to identify putative microglia (CD45^mid^CD11b^+^), macrophage/activated microglia (CD45^hi^CD11b^+^), and lymphocyte (CD45^+^CD11b^–^) populations. CD45^+^CD11b^–^ cells were further gated for CD4 and CD8 expression. (C) Representative immunostaining of DG, CA3 and CA1 of hippocampi from Tmem119 ^GFP/+^Ccr2^RFP/+^ reporter mice in the hippocampi of Mock-infected mice at 7 (left) and 25 (right) dpi. Data are representative of one independent experiment.

**Figure S2:**
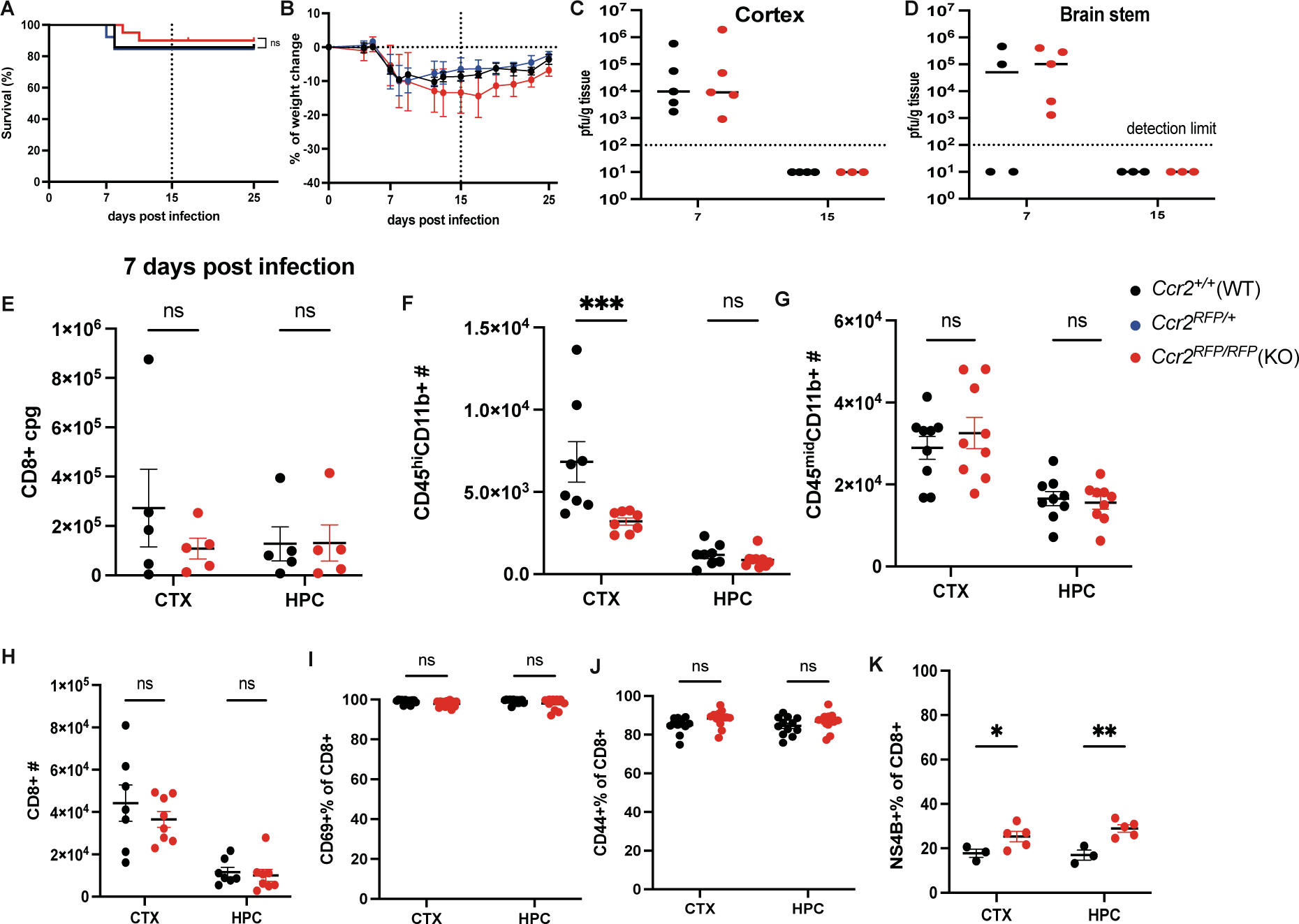
CCR2 is not required for the recruitment during acute infection or maintenance of CD8 T Cells post-WNV Infection. (A-D) Characterization of CCR2 KO mice post-WNV infection. Survival curve (A), weight loss changes (B) and viral burden at 7 or 15 dpi measured by plaque assay in the cortex (C) and brain stem (D) in WNV-infected Ccr2^+/+^ (WT), Ccr2^RFP/+^ and Ccr2^RFP/RFP^ (KO) mice. (E) Flow cytometry analysis showed no significant differences in the CD8 T cell counts per gram tissue in the CTX and HPC between WT and KO mice at 7 dpi. (n = 5 (WT), 5 (KO) mice per group) ns, not significant. (F-H) Cell counts of CD45^hi^CD11b^+^, CD45^mid^CD11b^+^ and CD8^+^ cells isolated from CTX and HPC of Ccr2^RFP/RFP^ and WT mice at 25 dpi. The number of CD45^hi^CD11b^+^ was decreased due to CCR2 deficiency. (I-J) Quantification of the percentage of CD8 T cells that are CD69^+^ (I) or CD44^+^ (J) from CTX and HPC of Ccr2^RFP/RFP^ and WT mice at 25 dpi. (K) Quantification of the percentage of CD8 T cells that are NS4B^+^ (specific for WNV) (n = 3 (WT), 4 (KO) mice per group). Data are representative of one independent experiment from 3 repeats. For F-J, Data were pooled from two independent experiments (n = 8 (WT), 9 (KO) mice per group). For E-K, data represent the mean ± s.e.m. and were analyzed by two-way ANOVA and corrected for multiple comparisons. *P < 0.05, **P < 0.005, ***P<0.0005. ns, not significant.

**Figure S3:**
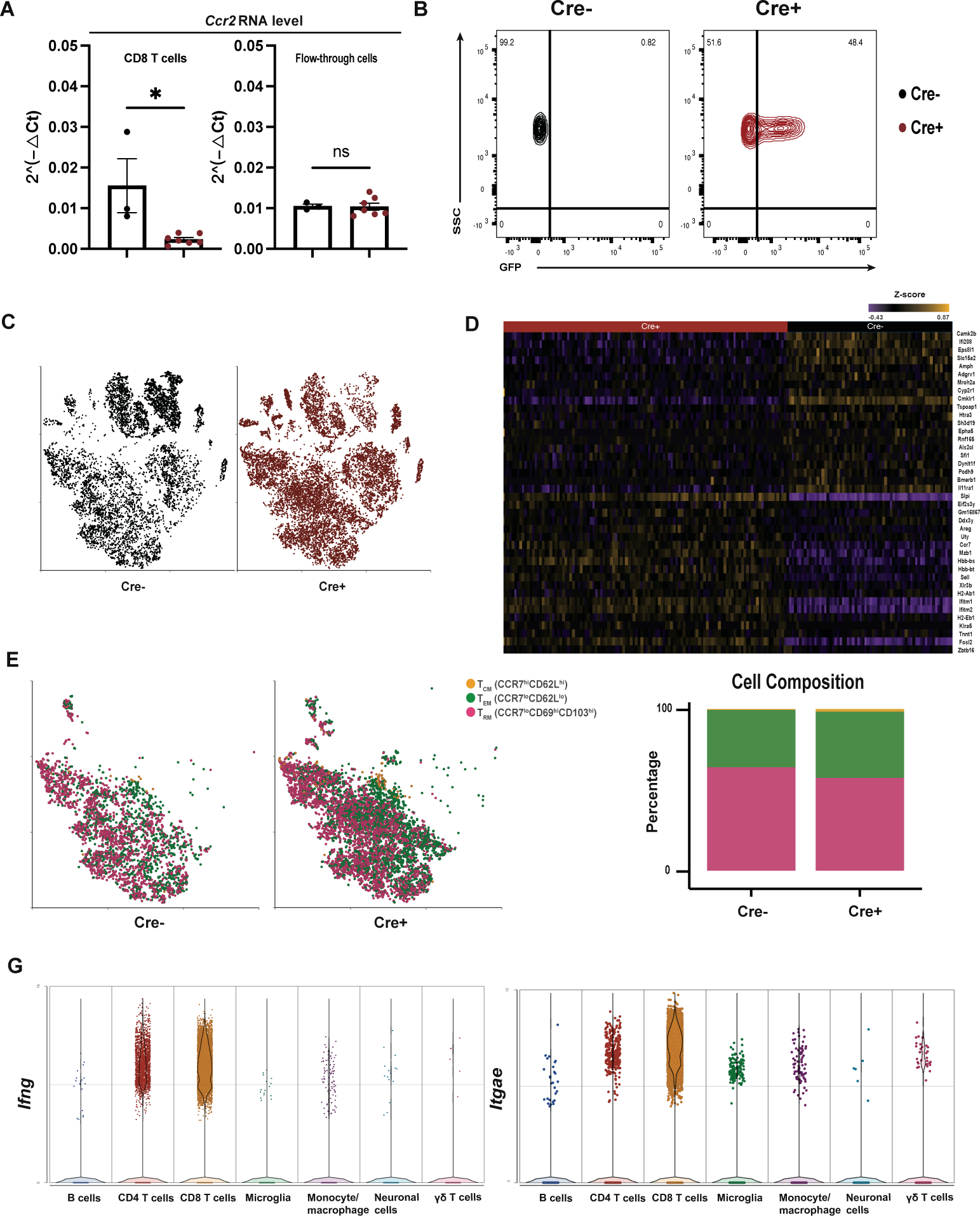
CCR2 expression is erased specifically in CD8 T cells from Cd8a^Cre+^Ccr2^fl/fl^ mice. (A) qRT-PCR validation of Ccr2 RNA depletion in CD8 T cells. CD8 T cells were magnetically isolated by negative selection from the forebrains of WNV-infected Cre^+^ and Cre^-^ animals at 30 dpi. Flow-through cells include microglia and other cell types from the CNS. Primers were designed to detect exon 3 of Ccr2 specifically. Statistical significance was calculated using an unpaired t-test. * P <0.05. (B) Flow cytometry analysis shows exclusive GFP expression on CD8 T cells in Cre^+^ mice. (C) t-SNE plots visualizing immune cells from the forebrains of WNV-infected Cre^+^ and Cre^-^ mice at 30 dpi. Clusters are colored by genotype. (D) Heatmap of DEGs comparing Cre^+^ and Cre^-^ samples. (E) t-SNE plots visualizing the CD8 T cell cluster from Cre^+^ and Cre^-^ mice. Clusters are colored by memory T cell types as shown in the legend. Central memory T cell (T_CM_, CCR7^hi^CD62L^hi^), effector memory T cell (T_EM_, CCR7^lo^CD62L^lo^), resident memory T cell (T_RM_, CCR7^lo^CD69^hi^CD103^hi^). (F) Proportion of each memory T cell type. T_RM_ was the major T cell type in the CNS at 30 dpi. (G) mRNA expression levels of Ifng and Itgae by each cell cluster. T cells were the main producer of Ifng post-WNV infection. Itgae was mostly expressed in the CD8 T cell cluster.

**Figure S4:**
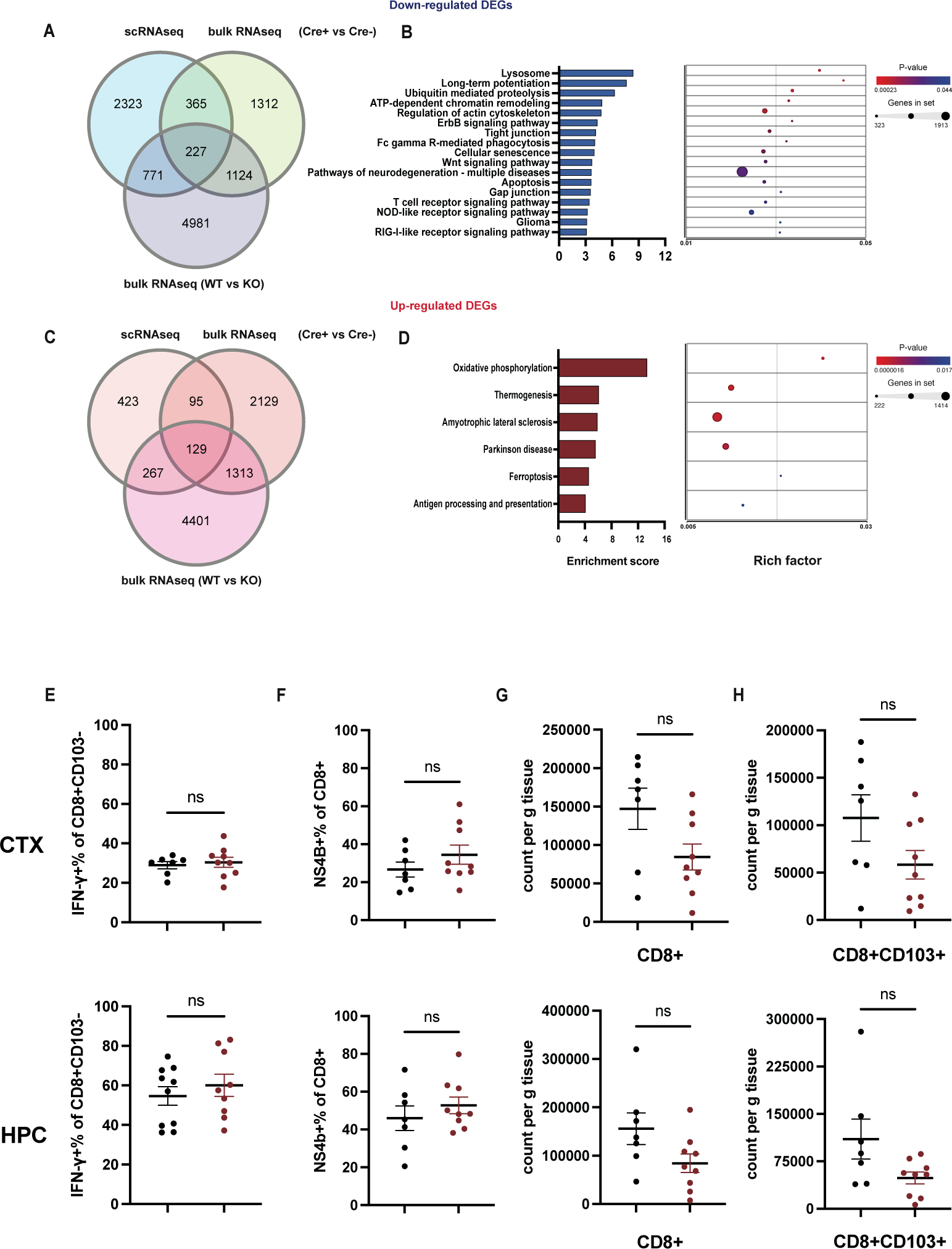
Characterization of Cd8a^Cre+^Ccr2^fl/fl^ mice. (A and C) Venn diagram of up- and down-regulated DEGs from bulk RNA-seq of WT and KO mice with those from scRNA-seq and bulk RNA-seq of Cre^+^ and Cre^-^ mice. (B and D) Biological pathways from GSEA based on overlapped DEGs from Fig. S4 A. (E) Quantification of the percentage of CD103^-^ CD8 T cells that are IFN-γ^+^. (F) Quantification of the percentage of CD8 T cells that are NS4B^+^. (G-H) Counts per gram CTX and HPC of CD8^+^ and CD8^+^CD103^+^ cells from Cre^+^ and Cre^-^ mice at 30 dpi. For E-H, data were pooled from two independent experiments (n = 7 (Cre^-^), 9 (Cre^+^) mice per group). Data represent the mean ± s.e.m. and were analyzed by unpaired t-tests. ns, not significant.

**Figure S5:**
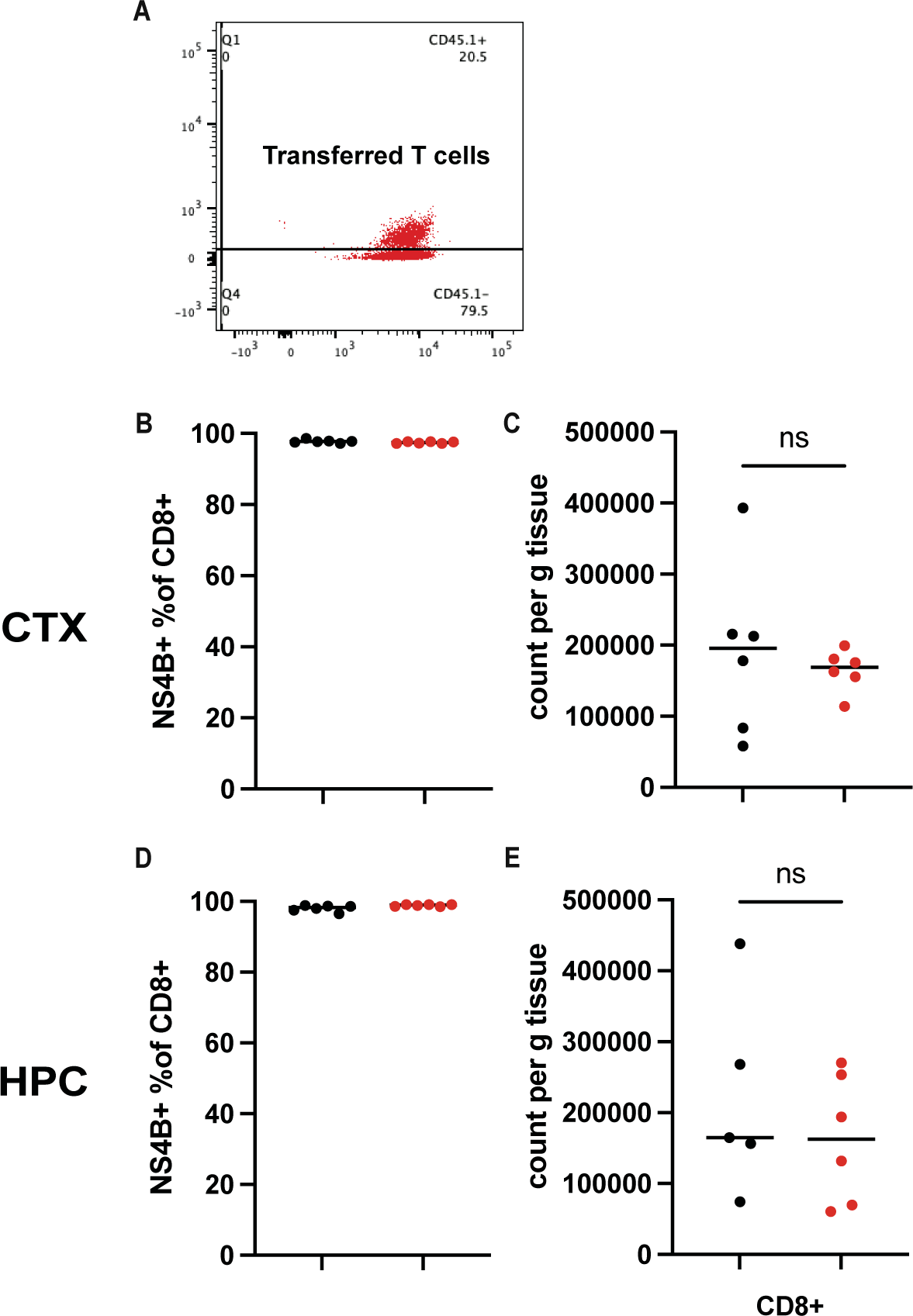
Adoptive transfer of CD8 T cells confirms that the retention of CD8 T cells in the CNS was not affected by CCR2 deficiency. (A) Representative dot plot of flow cytometry distinguishing between CD45.1^+^CD45.2^+^ transferred T cells and CD45.2^+^ cells from isolated CTX and HPC at 30 dpi. (B and D) All transferred CD8 T cells were NS4B^+^. (C and E) Counts per gram CTX and HPC of CD45.1^+^ CD8 cells from recipient mice at 30 dpi. Data are representative of one independent experiment from 3 repeats. Data represent the mean ± s.e.m. and were analyzed by unpaired t-tests (n = 6 (WT WNV-I), 6 (KO WNV-I) mice per group). *P < 0.05. ns, not significant.

